# Resting fMRI-guided TMS evokes subgenual anterior cingulate response in depression

**DOI:** 10.1101/2022.09.08.507012

**Authors:** Romain J. Duprat, Kristin A. Linn, Theodore D. Satterthwaite, Yvette I. Sheline, Ximo Liang, Gabriela Bagdon, Matthew W. Flounders, Heather Robinson, Michael Platt, Joseph Kable, Hannah Long, Morgan Scully, Joseph A. Deluisi, Michael Thase, Mario Cristancho, Justin Reber, Russell T. Shinohara, Desmond J. Oathes

## Abstract

Depression alleviation following treatment with repetitive transcranial magnetic stimulation (rTMS) tends to be more effective when TMS is targeted to cortical areas with high resting state functional connectivity (rsFC) with the subgenual anterior cingulate cortex (sgACC). However, it has not yet been confirmed that rsFC-guided TMS coil placement leads to TMS modulation of the sgACC. For each participant (N=115, 34 depressed patients), a peak rsFC cortical ‘hotspot’ for the sgACC and control targets were prospectively identified. Single pulses of TMS interleaved with fMRI readouts were then administered to these targets and established significant downstream fMRI BOLD responses in the sgACC. We then marked an association between TMS-evoked BOLD responses in the sgACC and rsFC between the stimulation site and sgACC. This effect was qualified by a difference between healthy and patient participants: only in depressed patients, positively connected sites of stimulation led to the strongest evoked responses in the sgACC. Our results highlight rsFC-based targeting as a viable strategy to causally modulate sgACC subcortical targets and further suggest that cortical sites with high positive rsFC to the sgACC might represent an alternative target for the treatment of depression.

## Introduction

High frequency (HF) rTMS has been FDA cleared as a therapy for treatment-resistant major depressive disorder (MDD) for over 10 years. The standard treatment protocol uses repeated trains of stimulation over the left dorsolateral prefrontal cortex (DLPFC), a region known for its role in top-down control and affective regulation known to be hypoactive in depressed populations ^1^. This region represents a large portion of the prefrontal cortex and is not well defined anatomically. Moreover, inter-individual variability limits the reliability of classic localization techniques used in clinical practice for treating depression with rTMS, such as the 5/6 cm rule (where the TMS coil is placed at fixed distance from TMS-evoked motor response ‘hotspot’ for all patients regardless of head size) or a coordinate based on scalp measurements (EEG F3). The use of more sophisticated neuronavigation devices allowing for precise anatomical identification of cortical targets may confer a clinical advantage ^2^, although the response rate to HF-rTMS in major depressive disorder (MDD) has not substantially improved since its introduction. The question remains as to where in the DLPFC (if in the DLPFC at all) stimulation would be the most clinically effective.

Resting state functional connectivity (rsFC) is a widely used method to assess functional links between brain regions or networks associated with psychopathology. Recently, several neuroimaging studies have highlighted the potential of rsFC as a predictor of clinical response to rTMS in depression. For example, rsFC between the site of stimulation and the subgenual anterior cingulate cortex (sgACC) before treatment was found to predict clinical efficacy ^3^. In another study of several potential Fmri predictors, it was found that only rsFC from the site of stimulation to the sgACC independently predicted treatment response ^4^.

The role of the sgACC in MDD pathophysiology has been highlighted by several neuroimaging studies showing hyperconnectivity ^5^ and abnormal rsFC in patients ^6,7^. In response to rTMS treatment, MDD patients exhibit changes in sgACC rsFC perhaps due to sgACC pathway engagement and plastic changes through this and closely related pathways ^8^. Taken together, these results indicate that targeting sgACC pathways might represent an effective brain stimulation protocol for treating depression. An innovative accelerated rTMS treatment based on connectivity with the sgACC has shown high response and remission rates in two studies of treatment resistant depressed patients ^9,10^.

However, the relevance of specific functional pathways connected to the site of stimulation for clinical efficacy is still poorly understood. The current sgACC rTMS depression literature focuses exclusively on rsFC targets anticorrelated with the sgACC ^3,11,12^. The meaning of anticorrelated fMRI signals is still a source of debate ^13^. Experiments combining rsFC with electrocorticography indicate that there are indeed negative and positive functional connections between distributed brain regions ^14^. However, when electrically stimulating areas of positive, null and negative functional connectivity, positively connected stimulation sites more effectively modulated downstream functionally connected regions ^15^. Although electrical brain stimulation and TMS are not identical, positively connected targets are worth exploring. This strategy is at apparent odds with clinical evidence in depression rTMS suggesting that stimulating more negatively correlated sites yields better clinical outcomes for treatment of patients ^3^. On the other hand, the association between negative sgACC-DLPFC connectivity and symptom improvement ^11^ does not directly speak to whether TMS engages this circuit, whether anticorrelation optimally targets the sgACC, or what brain changes contribute to clinical outcomes. TMS applied while functional brain activity is recorded can address the first two of these questions. TMS/fMRI can also be used broadly to probe brain circuit/pathway integrity in patient studies that include treatment ^16^.

In a small sample of healthy participants, we previously determined that a mix of sgACC-seeded positive and negative rsFC cortical sites, when stimulated with TMS, were successful at modulating (down regulating) the sgACC per our interleaved TMS/fMRI protocol ^17^. Evoked sgACC BOLD responses did not depend on pain ratings, somatomotor brain responses, cortical distance, or physiological arousal.

In the current replication and extension of our prior results, we endeavored to determine whether individually defined, positively connected sites of stimulation were effective in modulating the sgACC, critically evaluating whether rsFC at the site of stimulation was predictive of the amplitude of the sgACC evoked response (ER). In addition, we tested the relevance of TMS/fMRI to the study of depression by including a depressed patient sample and by looking at associations between symptom endorsements and the TMS-evoked sgACC fMRI BOLD response. Given previous findings in the electrical stimulation literature, we hypothesized that stimulating sites with high positive rsFC would be the most effective way of modulating sgACC activity. Per our prior work and the putative preferred direction of a negative ER (to counteract MDD sgACC hyperactivity), we predicted that applying single-pulse TMS to positively correlated sites in the prefrontal cortex would generate a stronger negative BOLD response in the sgACC compared to applying TMS to weakly or negatively correlated sites. We hypothesized that these effects would hold for depressed and healthy populations and that a more negative evoked sgACC BOLD response would be associated with relatively less severe symptoms of depression per better prefrontal cortex regulation of the sgACC.

## Materials / Subjects and Methods

### Participants

Participants (N=115) had ages ranging from 18 to 56 years (mean 29.53, standard deviation 9.86), had completed high school to doctoral education, had no history of a neurological or psychiatric condition (for healthy controls), and were not taking any psychoactive medications (including patients). The sample consisted of 46 male and 69 female participants, including 34 MDD patients and 81 healthy controls. During the TMS/fMRI session, participants received single-pulse TMS over their individualized rsFC-based target designed to modulate the sgACC as well as an active control target. Stimulation order was counterbalanced. Depending on participant and scanner availability, some healthy controls (N=17) returned for a second TMS/fMRI session with updated individualized targets. All participants gave written informed consent after receiving a complete description of the study according to the Declaration of Helsinki, and the study was approved by the University of Pennsylvania IRB.

### Baseline MRI session

The MRI data were acquired on a 3 Tesla Siemens Prisma scanner. For baseline scans (T1w and resting state) a standard 64 channel head coil was used (Erlangen, Germany), whereas for the interleaved TMS/fMRI scans a custom “birdcage” head coil was used to accommodate the MRI-compatible TMS coil (RAPID quad T/R single channel; Rimpar, Germany) and custom-built TMS coil holder. Two baseline multiband resting state fMRI scans were acquired with opposite phase encoding directions A>>P and P>>A (TR=800ms, TE=37ms, FA=52°, FOV=208mm, 2×2×2mm voxels, 72 interleaved axial slices with no gap, 420 volumes). Subjects were instructed to keep their eyes open and remain as still as possible while fixating on a central fixation cross. Structural images consisted of a high-resolution multi-echo T1-weighted MPR image (TR=2400ms, TI=1060ms, TE= 2.24ms, FA=8°, 0.8×0.8×0.8mm voxels, FOV=256mm, PAT mode GRAPPA, 208 slices).

### TMS/fMRI session

TMS within the MRI bore was delivered using an MRI-compatible Magventure MRI-B91 air-cooled TMS coil connected to a Magpro X100 stimulator (Magventure Farum, Denmark). The TMS coil was positioned within the MRI head coil with the cable end running through the MRI bore for a posterior-to-anterior induced current. Resting motor threshold (rMT) was obtained by visual observation of motor activity in the abductor pollicis brevis of the right hand in the MRI room to account for filters, cable length and any magnetic field interference. Stimulation intensity was then set to 120% motor threshold for all single-pulse stimulation. The gating of MRI volume acquisition (TR=2000ms, TE=30ms, FA=75°, FOV=192mm, 3×3×4mm voxels, 32 interleaved axial slices, 174 volumes) and TMS pulses were initiated by TTL triggers sent through a parallel port with E-prime 2.0 (Psychology Software Tools, Sharpsburg Pennsylvania USA) installed on a Windows PC. Between each volume acquisition, a gap of 400ms was inserted to allow single-pulse TMS to be delivered at 200ms without contaminating the subsequent volume.

Single pulses were delivered in 12 mini-blocks of 7 stimulations separated by 2400ms (1TR and the gap) and included 0, 1, or 2 ‘catch trials’ (random order and block), during which spacing was separated by an extra 2400ms with no TMS pulse to avoid easy prediction of TMS delivery by participants. The mini-blocks were themselves separated by 7 TRs, and 71 total stimulations were given per site over 174 volume acquisitions. Sequences where participants were not able to provide low-motion scans (relative motion >0.2mm) were excluded from the following analyses (26/275).

### MRI processing

MRI data were preprocessed using the eXtensible Connectivity Pipelines (XCP) developed at the PennLINC, University of Pennsylvania ^18^ to minimize head motion artifacts in rsFC data. ANTS cortical thickness ^19^ was used for segmentation of the structural T1w image and to compute transforms between the high-resolution anatomical image and the MNI template using the SyN algorithm.

For rsFC maps (i.e. baseline resting state), the first 2 volumes were discarded for scanner equilibration, and fMRI volumes were realigned to the volume with the least motion. Skull stripping was done using FSL’s BET, and AFNI’s 3dDespike was used to identify and interpolate outliers in the time series. Images were then demeaned and detrended (linear and polynomial detrend). Confound models were computed for grey matter, white matter, cerebrospinal fluid, relative motion, and global signal ^20^. Band-pass filtering was applied at 0.01-0.08 Hz.

Motion artifact in task-free data was modelled as a linear combination of 36 time series ^20^ and removed from the BOLD signal using a general linear model to estimate residuals. To ensure that the spectral content of the BOLD timeseries and nuisance regressors matched, temporal filtering of data was performed simultaneously with confound regression ^21^.

Baseline functional connectivity values used for targeting were computed from our sgACC ROI seed, a sphere of 1cm diameter at MNI coordinates (−2, 18, -8) transformed into individual space, representing abnormalities in depression ^8^ in the left hemisphere ipsilateral to stimulation, as in our prior work ^17^.

For the ER (interleaved TMS-fMRI single-pulse probe), the analysis was carried out using the FMRI Expert Analysis Tool (FEAT, v6.00). Brain extraction was performed on both structural and functional images with the Brain Extraction Tool (BET; (21)). The EPI time series were motion corrected through FSL MCFLIRT using the six standard motion regressors. High-pass filtering was also applied to the images with a cut off of 100, and spatial smoothing was applied with a 5 mm kernel (FWHM).

Each TMS pulse was considered an instantaneous event (effective TR of 2400ms due to the 400ms break needed for stimulation between volume acquisitions, protocol TR of 2000ms) and convolved with a gamma-shaped hemodynamic response function. After model estimation, the resulting maps of the parameter estimates and contrast values of the parameter estimates were converted into percent signal change (TMS on vs TMS off) to quantify the ER. Registration to the MNI template was accomplished by applying the non-linear transforms computed by ANTs to warp maps from subject native space to the MNI template coordinate space.

### TMS target design

The stereotaxic neuronavigation system from Brainsight (Rogue Research, Montreal, Quebec, Canada) was used to localize targets anatomically and functionally. Individualized sgACC targets were chosen as follows: the rsFC maps were registered and loaded into individual space (Brainsight), and then the highest correlation value for the sgACC seed, preferentially located in prefrontal cortex, was chosen as the stimulation site (Figure 1A). As control sites, we used sites designed for parallel studies, such as the motor hand knob and/or amygdala rsFC individualized sites—i.e., sites not chosen for their rsFC to the sgACC—that included a variety of distributed cortical sites with varying levels and valence of rsFC with the sgACC (Figure 1). Each target was localized using Brainsight following the shortest trajectory from the cortical target perpendicular to the scalp and marked on an individual Lycra swim cap that the participant wore during scanning.

**Figure 1:**
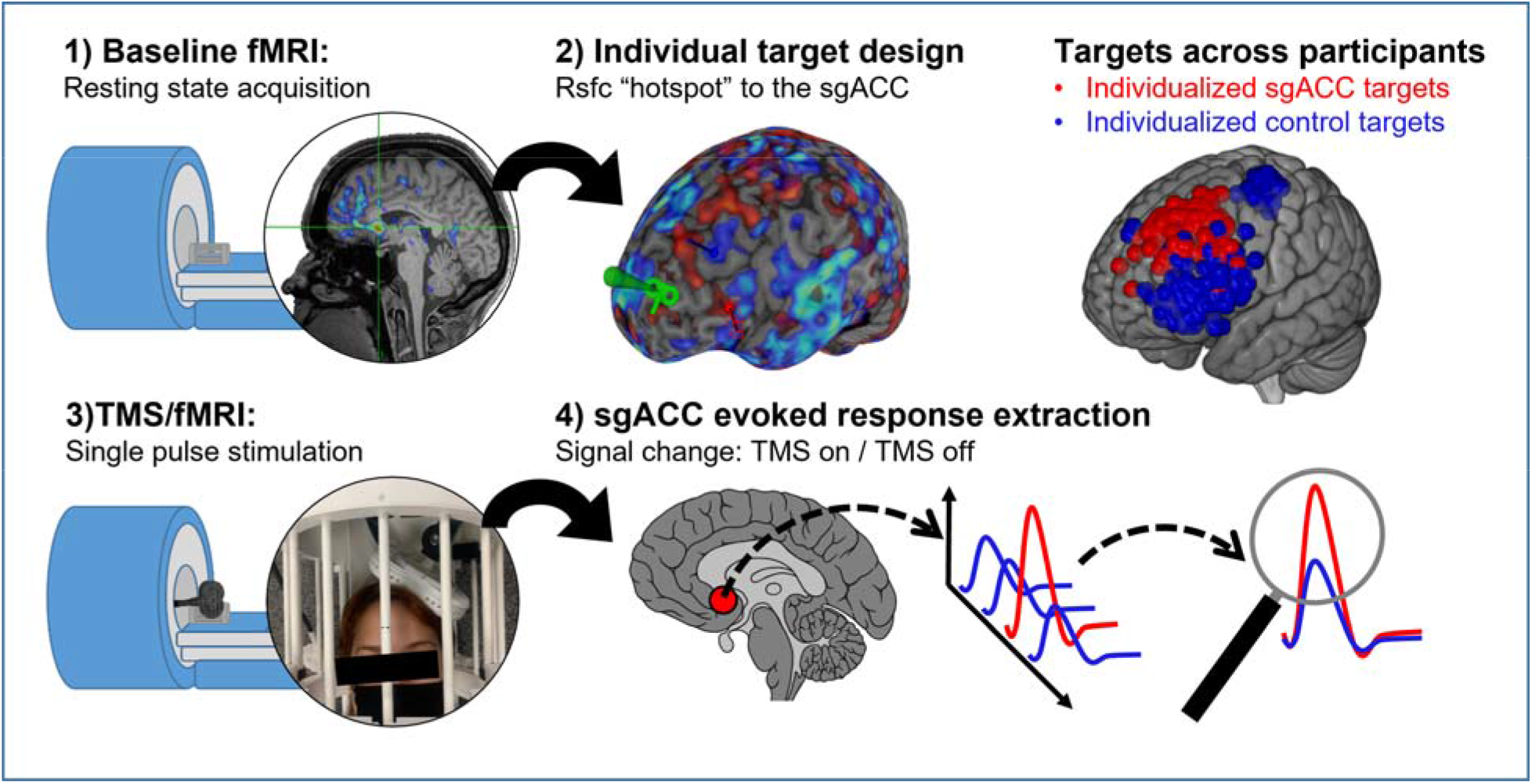
Experimental protocol. Representation of the main steps of the experimental protocol. In the top right corner, sites of stimulation across participants are overlaid on the MNI brain template. Red spheres represent Individualized sgACC targets (rsFC hotspot to the sgACC) and blue spheres represent individualized control targets (anatomical landmark or rsFC hotspot to a different seed). The sagittal section drawing has been modified from Patrick J. Lynch “Brain human sagittal section” ^1^.

### Statistical analysis

First, to test for sgACC engagement to stimulation of the individualized sgACC rsFC sites, we used a two tailed one sample t-test to test that the mean of the average sgACC ER (percent signal change for the contrast between TMS on vs TMS off averaged within the sgACC) was different from zero following single-pulse stimulations of the individualized targets. For the few participants (N=17) that underwent two TMS/fMRI sessions and thus had two individualized sgACC targets, we retained for analysis the rsFC-based target with highest positive rsFC to the sgACC.

Second, we tested whether rsFC at any site of stimulation was associated with the TMS ER in the sgACC, pooling data from all sites (individualized sgACC and control sites in MDD and healthy control subjects) and including both observations from the participants who underwent two TMS/fMRI sessions. To account for within-subject correlation among repeated measures in the analysis of fMRI BOLD ER extracted from our sgACC region of interest (ROI), we used generalized estimating equations (GEE). The analysis was run in “R” *(R Core Team, 2020)* version 3.6.3 with the “geepack” package ^22^. We fit a GEE model with the ER extracted from the sgACC as the response variable and rsFC between the sgACC seed and site of stimulation, group (MDD vs HC), and their interaction as explanatory factors. We refer to this first GEE model as the unadjusted GEE analysis. To determine whether the effects were robust to potential confounding, we then fit an adjusted model using GEE that additionally included the following variables: age, gender, education level, scalp discomfort (self-rated intensity of pain), TMS dose (distance adjusted proportional to motor hotspot and threshold), and order (sequence in which the targets were stimulated within sessions). We refer to this second GEE model as the adjusted GEE analysis. Only cases with completely observed covariates were included in the adjusted analysis due to a small percentage of incomplete cases (6/115 subjects). We used an exchangeable correlation structure for the main analyses but also re-fit the models using an independent correlation structure as a sensitivity analysis (supplementary tables 1 and 2). To visualize the association between rsFC and sgACC ER among the rsFC individualized sites, we plotted the adjusted marginal effect for each group over the observed data and reported the resulting intercepts and slopes of the linear effect by group.

Finally, to investigate whether the relative magnitude of sgACC ER following stimulation to individualized targets was related to depression severity and affect in our MDD group, we performed exploratory correlation analyses with the Positive and Negative Affect Schedule (PANAS) ^23^ and depression subscores of the Depression Anxiety Stress Score (DASS21) ^24^.

## Results

### Individualized targets engage the sgACC

With fMRI-guided individualized sgACC-seeded positive FC cortical targets, we replicated our prior work demonstrating an average negative TMS-evoked BOLD response in the sgACC across the combined sample (M=-0.24) t = -6.5, df = 112, p<0.001, d = 0.61; healthy participants alone (M=-0.23) t = -4.95, df = 80, p<0.001, d = 0.55; and patients alone (M=-0.28) t = -4.46, df = 32, p<0.001, d = 0.78 by one-sample t-tests (vs. 0).

### rsFC at the site of stimulation is associated with TMS-evoked fMRI BOLD in the sgACC

The unadjusted GEE analysis showed a significant interaction between rsFC and group (MDD/HC), PE = 0.6; SE=0.3; W=3.85; p=0.049 for the sgACC ER, driven by the negative relationship between rsFC and ER in the sgACC being stronger in the patient group than in the healthy control group. After adjusting for potential confounders, the interaction effect was similar in magnitude and remained significant: PE = 0.505; SE=0.254; W=3.95; p=0.047 (see Table 1).

**Table 1:**
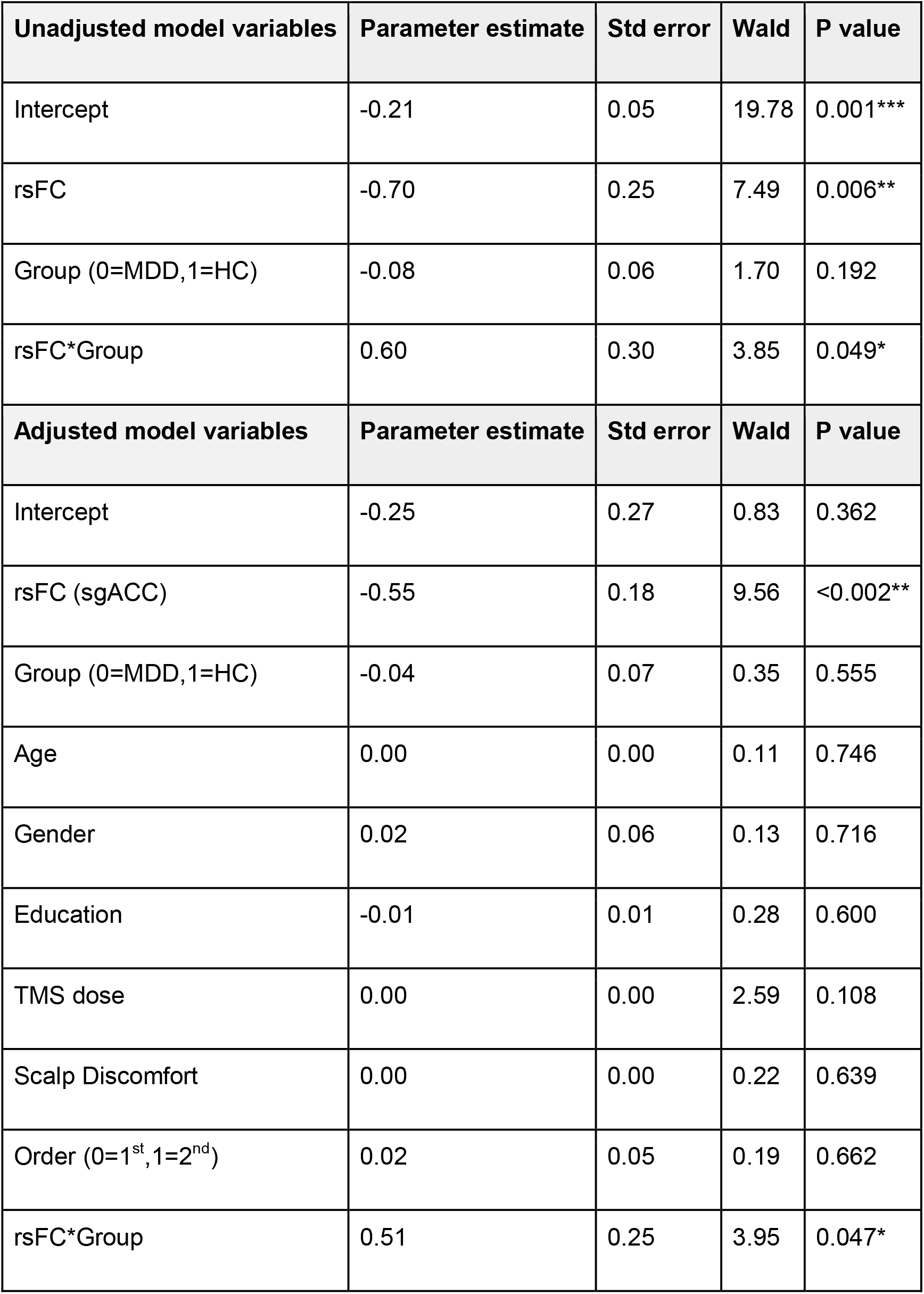
Gee Models. Unadjusted GEE model (N=115) with exchangeable correlation structure (_α_ = -0.002) and adjusted GEE model (N=109) with exchangeable correlation structure (_α_ = 0.03). Significance code: <0.001 ‘***’<0.01 ‘**’<0.05 ‘*’.

| **Unadjusted model variables** | **Parameter estimate** | **Std error** | **Wald** | **P value** |
| --- | --- | --- | --- | --- |
| Intercept | -0.21 | 0.05 | 19.78 | 0.001*** |
| rsFC | -0.70 | 0.25 | 7.49 | 0.006** |
| Group (0=MDD,1=HC) | -0.08 | 0.06 | 1.70 | 0.192 |
| rsFC*Group | 0.60 | 0.30 | 3.85 | 0.049* |
| **Adjusted model variables** | **Parameter estimate** | **Std error** | **Wald** | **P value** |
| Intercept | -0.25 | 0.27 | 0.83 | 0.362 |
| rsFC (sgACC) | -0.55 | 0.18 | 9.56 | <0.002** |
| Group (0=MDD,1=HC) | -0.04 | 0.07 | 0.35 | 0.555 |
| Age | 0.00 | 0.00 | 0.11 | 0.746 |
| Gender | 0.02 | 0.06 | 0.13 | 0.716 |
| Education | -0.01 | 0.01 | 0.28 | 0.600 |
| TMS dose | 0.00 | 0.00 | 2.59 | 0.108 |
| Scalp Discomfort | 0.00 | 0.00 | 0.22 | 0.639 |
| Order (0=1<sup>st</sup>,1=2<sup>nd</sup>) | 0.02 | 0.05 | 0.19 | 0.662 |
| rsFC*Group | 0.51 | 0.25 | 3.95 | 0.047* |

Based on the estimated marginal effect of rsFC in MDD, a 0.1 increase in resting state correlation between the site of stimulation and the sgACC would be associated with a decrease of 6.9% (stronger negative BOLD) in the sgACC ER (see Figure 2). As seen in the scatterplot, for HCs, the average sgACC ER was negative, similar to our prior paper ^17^, but rsFC valence was not associated with magnitude or direction of the sgACC ER. For MDD patients, on the other hand, the strongest modulations of activity (negative ER) in the sgACC were measured after stimulations over high positive rsFC targets.

**Figure 2:**
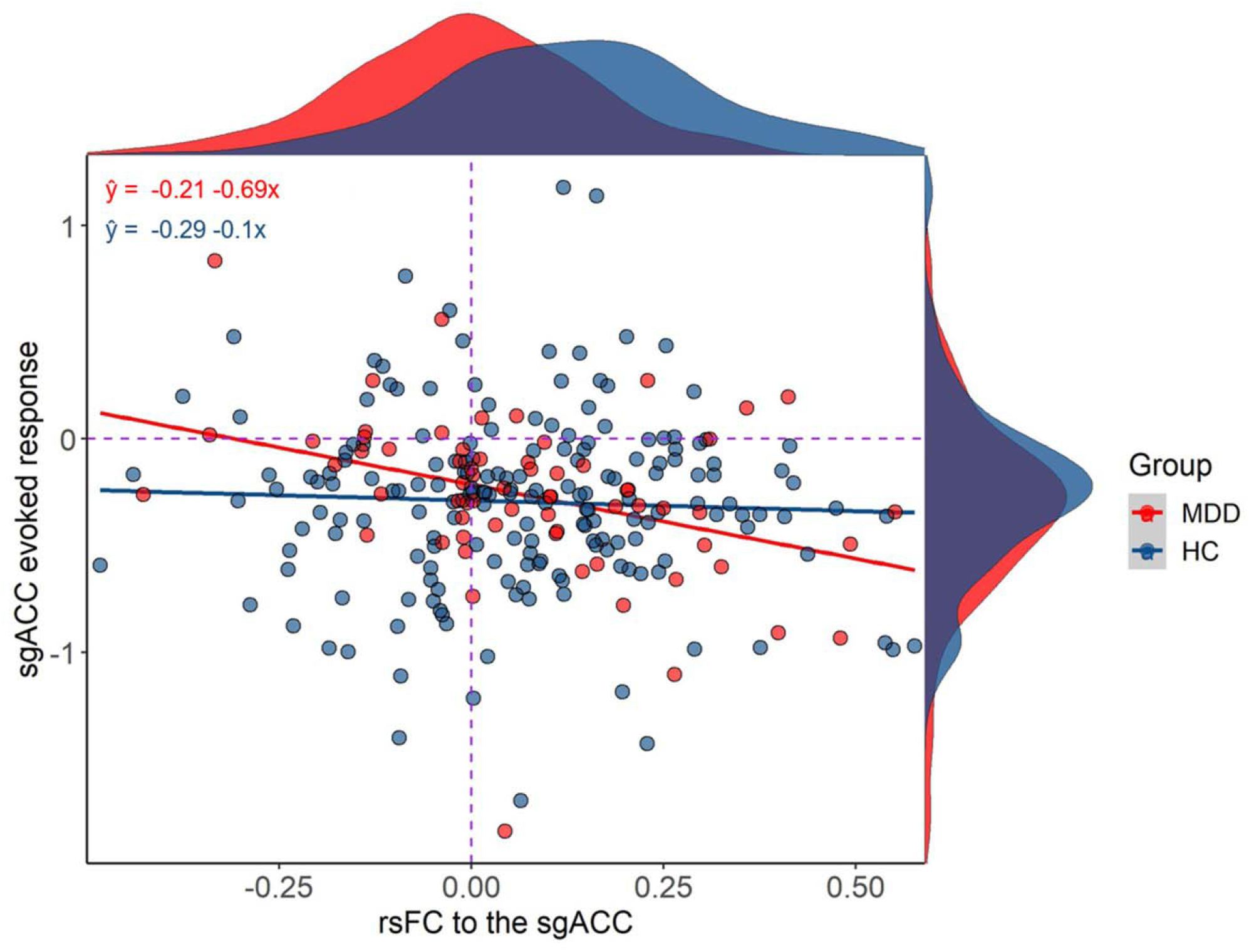
Evoked response as a function of function of connectivity. Marginal distributions and relationship between rsFC to the sgACC at the site of stimulation and ER in the sgACC for healthy controls (N=81) and MDD patients (N=34). The red and blue lines represent the within-group relationships estimated from the unadjusted GEE model with rsFC by group (MDD and HC) interaction

### sgACC ER amplitude and behavioral measures

In depressed patients, we explored the relationship between the sgACC ER and patients’ negative affect, positive affect, and DASS21 depression subscores. Based on the Pearson correlation, there was a significant positive association between the sgACC ER and the negative affect PANAS subscore in MDD (Figure 3): r = 0.44, t = 2.620, df = 29, p-value = 0.014 that survived Bonferroni correction across the three tests (p = 0.05/3 = 0.017). This association suggests a possible relevance of TMS-evoked BOLD responses in the sgACC to depression symptoms in patients.

**Figure 3:**
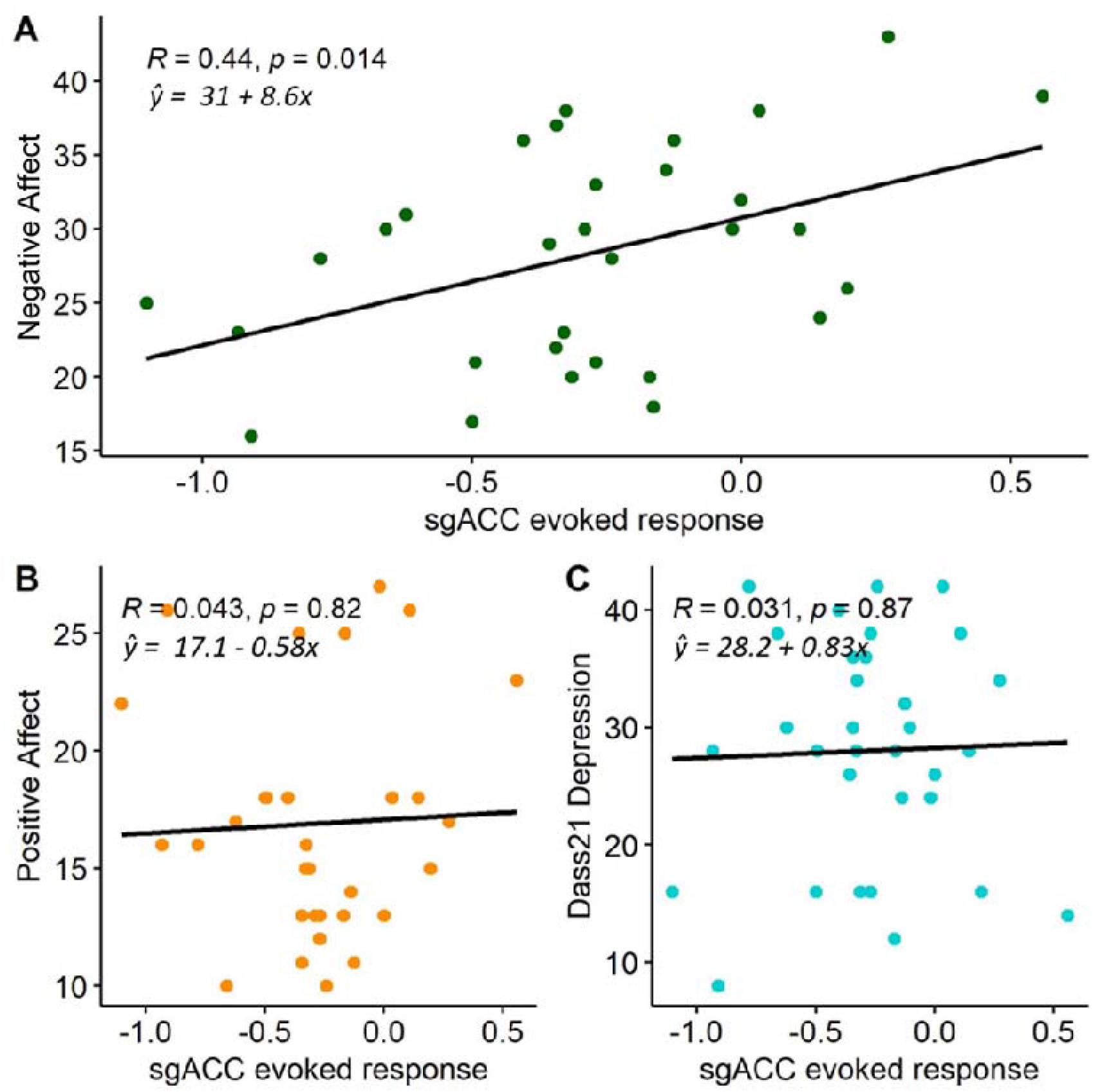
Relationship between behavioral measures and evoked response. Relationship between sgACC ER and (A) baseline Positive affect, (B) Negative affect, and (C) DASS21 depression subscores in MDD patients. The black lines represent the slope of a linear regression fit to the data.

No other association approached significance (Figure 3B/C).

## Discussion

With recent evidence that TMS treatment targeting sgACC functionally connected targets is more effective in treating depression than any prior published protocol, the importance of better understanding how functional connectivity at the site of stimulation relates to TMS-evoked brain activity changes is underscored. Our group has recently demonstrated that TMS can indeed effectively modulate the sgACC when applied to functionally connected cortical regions indexed by interleaved TMS/fMRI ^17^. In the present study, we replicated this finding in a large independent cohort of healthy controls as well as extended the finding to a depressed patient cohort. In addition, we determined that the amplitude and valence of functional connectivity between the sgACC and site of stimulation were critical in determining the TMS evoked sgACC response, especially in depressed patients. Interestingly we also found that baseline self-reported negative affect was associated with TMS-evoked brain responses among patients.

In support of the focus on sgACC pathways for TMS, our results provide evidence of a depression-sensitive causal pathway between the prefrontal cortex and sgACC ^17^. This pathway has been previously hypothesized based on neuroimaging studies of rTMS-induced functional connectivity changes in MDD (29,30). In our MDD group, stimulation of high positive rsFC sites was especially effective in modulating the sgACC, and rsFC at the stimulation site predicted engagement of the sgACC with TMS. Given the known association between baseline negative rsFC at the site of stimulation and clinical outcomes observed in several studies ^3,8^, these results suggest that stimulation over the anticorrelated circuit might work though a different mechanism, such as gradually strengthening an initially weak connection to the sgACC and/or an associated network in patients rather than robustly initially modulating the sgACC. Both fMRI and EEG studies ^3,25^ suggest that the sgACC can be modulated as the consequence of normalizing network connections. Whether a more direct modulation of the sgACC by stimulating high positive rsFC sites would result in better or quicker clinical outcome is unknown and requires further investigation. The induced response from single-pulse stimulation is likely related to, but not directly testing, an rTMS treatment outcome, but nevertheless it can offer insights into the potential of a pathway to engage a distant region. Our results support the relevance of a prefrontal-sgACC circuit to negative affect, in that lower reported negative affect at baseline was associated with stronger TMS modulation of the sgACC in patients.

Indirect synaptic connections resulting in inhibition of the sgACC could explain the negative signal change induced in our region of interest, which is in line with the assumed inhibitory role of the prefrontal cortex over affective regions ^26^. Future research incorporating electrophysiology would be useful to substantiate the neuronal process underlying our negative evoked BOLD responses. Nevertheless, our results suggest that positive rsFC may be an especially fruitful indicator of how artificially stimulated pathways respond to TMS in patients ^14^ and that individual rsFC-based targeting is effective as a method to guide cortical stimulation to affect subcortical neural responses.

This study has limitations. First, these initial TMS/fMRI mappings do not include an rTMS treatment protocol comparing the sites of stimulation. rTMS treatment mechanism(s) of action might rely on different neural circuit effect(s) disparate from those included in our analysis (i.e., single-pulse ERs). Second, our depressed patients were not selected for treatment resistance which, at the moment, is the population approved by the FDA to receive rTMS treatment and hence is the most studied population in rTMS studies. Our patients were also unmedicated, which yields a cleaner brain response but limits the generalization of our findings to typical depressed clinical populations who are often medicated. Finally, we collected fewer control sites with strong negative connectivity, which somewhat limits our ability to extrapolate our results to rsFC outside of our sampled range.

Nevertheless, this study shows that using baseline rsFC is a useful approach for targeting and modulating distant brain regions such as the sgACC. On average, stimulation of a prefrontal cortical site with maximum functional connectivity to the sgACC results in an acute modulation of activity in that region. Our results suggest that in MDD patients, sites of high positive functional connectivity yield stronger, perhaps more direct connections to the sgACC. Consequently, our study supports a proposition that cortical sites with high positive rsFC to the sgACC might represent an especially promising new target for the treatment of MDD. More neuroimaging data, especially interleaved TMS/fMRI of this circuit in depressed patient samples preceding and following an rTMS intervention over such sites are needed to better understand the link between rsFC, TMS-evoked brain responses, and modulation of the sgACC as it relates to clinical response.

## Authors contributions

R.D., H.L., M.S., J.D., G.B., M.F. and H.R. conducted the experiment; D.O., T.S., Y.S., M.P., J.K. designed the experiment; R.D., T.S., K.L., X.L., R.S., D.O., M.T., M.C., and J.R. analyzed the data and wrote the paper.

## Acknowledgements

This work was supported by the Penn Translational Neuroscience Initiative (TNI) 827636 (YIS, MP, DJO, JK), NIH R01 MH111886 (DJO), NIH RF1 MH116920 (DJO & TDS.), a NARSAD BBRF Young Investigator Award (DJO), and UO1 MH109991-03: “Dimensional connectomics of anxious misery (YIS).

## Materials & Correspondence

Desmond Oathes, Ph.D. Richards Biomedical Building, D306, 3700 Hamilton Walk, Philadelphia, PA 19104. Phone: (215) 573-9390. Fax: (215) 573-8556.

## Data Availability

The data that support the findings of this study are available from the corresponding author, upon reasonable request.

## Footnotes

1 Patrick J. Lynch, medical illustrator; C. Carl Jaffe, MD, cardiologist. http://patricklynch.net/https://creativecommons.org/licenses/by/2.5/

