## Supplemental information for "Resting fMRI-guided TMS evokes subgenual anterior cingulate response in depression"

**Supplementary Materials**

-Control sites

As control for the individualized sgACC targets we used previously designed sites not picked for their rsFC to the sgACC. Those controls sites included individual motor hand knob (site leading to visual motor responses in the abductor pollicis brevis of the right hand) and rsFC based sites designed around the left basolateral amygdala (BLA) rsFC hence individualized BLA targets. Individualized BLA targets design was similar to individualized sgACC targets i.e., the connectivity “hotspot” to the BLA was selected.

GEE sensitivity analysis

To test our models under more stringent conditions we re-fit the models using an independent correlation structure:

| **Model variables** | **Parameter estimate** | **Std error** | **Wald** | **P value** |
| --- | --- | --- | --- | --- |
| Intercept | -0.213 | 0.048 | 19.76 | <0.001*** |
| rsFC | -0.695 | 0.254 | 7.48 | 0.006** |
| Group (0=MDD,1=HC) | -0.075 | 0.06 | 1.7 | 0.192 |
| rsFC*Group | 0.596 | 0.304 | 3.85 | 0.049* |

Supplementary Table 1: Unadjusted GEE model (N=115) with independent correlation matrix. Significance code: <0.001 ‘***’<0.01 ‘**’<0.05 ‘*’.

| **Model variables** | **Parameter estimate** | **Std error** | **Wald** | **P value** |
| --- | --- | --- | --- | --- |
| Intercept | -0.26 | 0.27 | 0.93 | 0.336 |
| rsFC (sgACC) | -0.55 | 0.18 | 9.56 | 0.002** |
| Group (0=MDD,1=HC) | -0.04 | 0.06 | 0.37 | 0.545 |
| Age | <-0.01 | <0.01 | 0.08 | 0.778 |
| Gender | 0.02 | 0.06 | 0.13 | 0.717 |
| Education | <-0.01 | 0.01 | 0.23 | 0.633 |
| TMS dose | <0.01 | <0.01 | 2,66 | 0.103 |
| Scalp Discomfort | <0.01 | <0.01 | 0.19 | 0.667 |
| Order (0=1st,1=2nd) | 0.02 | 0.05 | 0.18 | 0.670 |
| rsFC*Group | 0.50 | 0.25 | 3.84 | 0.050* |

Supplementary Table 2: Adjusted GEE model (N=109) with independent correlation matrix. Significance code: <0.001 ‘***’<0.01 ‘**’<0.05 ‘*’.
